# Surface-sourced eDNA to track deep-diver cetaceans: the study case of Cuvier’s beaked whale (*Ziphius cavirostris*) at Caprera Canyon and surrounding areas (Western Mediterranean Sea)

**DOI:** 10.1101/2023.11.07.565978

**Authors:** G. Boldrocchi, L. Conte, P. Galli, R. Bettinetti, E. Valsecchi

## Abstract

Among cetaceans, the Cuvier’s beaked whale is considered an extreme diver, very challenging to be studied with standard monitoring methods due to its elusive behaviour and a preference for deep offshore waters. These limitations seem to be mitigated by the use of new molecular methodologies capable of intercepting small traces of DNA left in the environment (eDNA) by marine organisms. Moreover, the collection of water from the superficial layer of the sea represents a perfect case study for the targeting of marine mammals, as the constraints imposed by their nature implies periodic and frequent surfacing. Therefore, we designed and tested a taxon-specific primer set to infer the Cuvier’s beaked whale presence, with the aims of 1) examining the effectiveness of the eDNA technique to detect the presence of deep-diving cetacean in open waters, using the Cuvier’s beaked whale as case study; 2) providing data on the spatiotemporal occurrence of this species within the Canyon of Caprera; and 3) assessing the species occurrence in central northern Mediterranean Sea based on molecular traces. Results from this study demonstrated that collection of superficial waters is a valid approach to monitor deep-diving cetacean species, without the need for complementary visual survey monitoring. Specifically, this study provides evidence of the regular presence of the Cuvier’s beaked whale in the Canyon of Caprera, with a preference for bathymetry in the range of 700-1000 m. This study also showed a potential inshore movement of this species during fall, which is possibly related to migration of its cephalopod prey or a shift in prey preferences. Overall, the data presented here are particularly relevant in the optic of proposing the Caprera Canyon as an *Important Marine Mammal Area* (IMMA) in the Mediterranean Sea. At a wider level, this study also showed that the stronger positive signals were recorded in sampling stations located on submarine canyon systems, demonstrating the importance of these areas as elective habitats for the Cuvier’s beaked whale, thus the pivotal priority to their conservation.

## 1. Introduction

The identification of areas critical for species survival is essential for effective wildlife conservation, particularly when focusing on species of concern (Ambal et al., 2012; Stokes et al., 2015; Valsecchi et al., 2023). Threatened marine vertebrates are challenging to study as they are often rare or elusive, resulting in insufficient knowledge on their occurrence and distribution, which impedes management and limits effective conservation (Boldrocchi and Storai, 2021; Kiszka et al., 2007; Smith et al., 2021). These limitations seem to be somewhat mitigated by the use of new molecular methodologies capable of intercepting small traces of DNA left in the environment (eDNA) by marine organisms (e.g. Bohmann et al., 2014). This approach is also advantageous because it is suitable for involving citizen scientists who can easily be engaged in the sampling phase (Biggs et al., 2015), which simply consists of the collection of seawater samples (Agersnap et al., 2022), usually from the surface to facilitate the collection operation even for non-experts using simple and readily available equipment (Valsecchi et al., 2023). Moreover, the collection of water from the superficial layer of the sea does not appear to represent a limitation when targeting marine mammals, as the constraints imposed by their nature implies periodic and frequent surfacing in order to breathe. In fact, the first researches that have used eDNA for the study of marine mammals involved the collection of seawater samples from the “footprint” left behind when the animals break through the water surface (Alter et al., 2022; Baker et al., 2018; Székely et al., 2021). However, while marine mammals represent a perfect target for superficial eDNA samplings, recent studies have also shown that the molecular signals degrade quickly after the animal submerges: bowhead whale DNA dropped by ∼4.5-fold 10 min after a dive (Székely et al., 2021), while for other cetacean species it was observed a decline in eDNA over the 15-min to 30-min intervals following the sighting (Alter et al., 2022). This can considerably reduce the possibility of molecularly identifying the passage of those marine mammals that spend short time on the surface, such as those considered “deep divers” like the sperm whales and other odontocetes belonging to the Ziphiidae family.

The research on those cetaceans considered deep-diving species is still particularly difficult as traditional monitoring surveys are often not very effective as these animals are rarely encountered and live in offshore habitats (Breck, 2006; Robbins et al., 2022). The use of species-specific qPCR assays can enhance the efficiency of detecting the presence of the eDNA of single target-species as this approach is more sensitive than the multispecies PCR approach (metabarcoding) targeting at broad taxonomic groups (e.g. Neice and McRae, 2021; Plante et al., 2021; Valsecchi et al., 2022).

Among deep divers, the Cuvier’s beaked whale (*Ziphius cavirostris*) is the only beaked whale species commonly found in the Mediterranean Sea, listed as Data Deficient on the IUCN Red List until recent years (Cañadas, 2012) when its status changed to Vulnerable (Cañadas and Notarbartolo di Sciara, 2018). This species is found both in the western and eastern basins of the Mediterranean Sea (Podestà et al., 2016). However, although the analysis of a massive long-term sightings dataset has recently better delineated the Mediterranean areas usually frequented by this species (Arcangeli et al., 2023; Gnone et al., 2023), information on the spatial ecology of the Cuvier’s beaked whale is still limited and mainly restricted to certain areas (such as the Pelagos Sanctuary for Mediterranean Marine Mammals and Alboran Sea, e.g. Moulins et al., 2007; Tenan et al., 2023; Tepsich et al., 2014; Torreblanca et al., 2022), leaving knowledge gaps where the study effort is non-existent (Cañadas and Notarbartolo di Sciara, 2018). Moreover, among cetaceans, the Cuvier’s beaked whale is considered an extreme diver (Figure 1), very challenging to be studied due to its elusive behaviour and a preference for deep offshore waters (Heyning, 1989; Tyack et al. 2006). The combination of all these traits results in a paucity of research and information compared to other cetaceans (Johnson et al., 2004; Schorr et al., 2014). Therefore, a comprehensive basin-wide survey and efficient alternative methodologies are increasingly necessary to fill essential knowledge gaps and to provide basic ecological information about presence and absence in those poorly surveyed areas (Piggott and Taylor, 2003). In this context, environmental DNA technique has been already proved to be fast and cheap methodology to detect rare and invasive species, including the elusive Mediterranean monk seal, one of the rarest pinnipeds at worldwide level (Valsecchi et al., 2023). The utility of eDNA is vast and not relying on visual observations, this approach overcome the cost of field monitoring, which might be expensive for offshore species like for the Cuvier’s beaked whale. Moreover, eDNA can spare time, biases, and avoid disturbance to animals associated with traditional monitoring methods (Jeunen et al., 2019; Thomsen and Willerslev, 2015).

**Figure 1.**
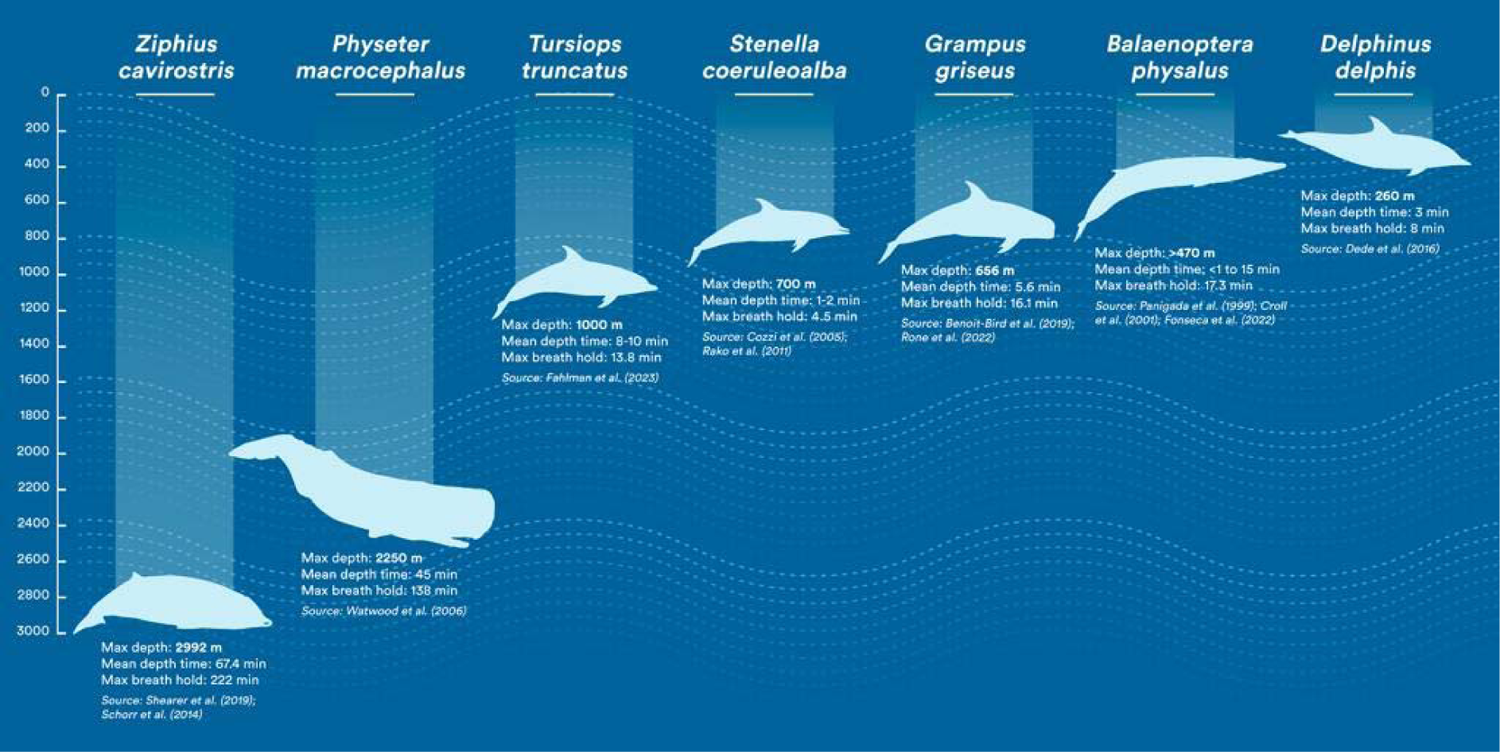
Comparison between cetacean species in their diving profiles according to literature records. Maximum reached depth, mean diving time and maximum breath hold are reported.

Considering the difficulties of monitoring the occurrence of Cuvier’s beaked whale, we designed and tested a taxon-specific primer set to infer the Cuvier’s beaked whale presence, with the following aims: 1) to examine the effectiveness of eDNA technique to detect the presence of deep-diving marine mammals in open waters, using the Cuvier’s beaked whale as case study; 2) to provide data, within the framework of *The Caprera Canyon Project* carried out by One Ocean Foundation, on the spatiotemporal occurrence of this species within the Canyon of Caprera, a potential important area for the species life-history (Bittau and Manconi, 2016); and 3) to assess the species occurrence in central northern Mediterranean Sea based on molecular traces. The results presented here are important not only for demonstrating the effectiveness of the eDNA approach for detecting deep-diving cetaceans (and thus all marine mammals) without visually detecting them, but also furthering our understanding of the Cuvier’s beaked whale geographic distribution and seasonality in the Mediterranean Sea, with a particular focus on the Canyon of Caprera, highlighting its importance for this elusive cetacean species. This information is particularly relevant in the optic of proposing the Caprera Canyon as an Important Marine Mammal Area (IMMA) within the basin.

## 2. Methods

### 2.1 Sampling Location

The study was carried out using three sets of samples. The first consisted of samples collected on a monthly basis, in three points along the rim of the Caprera Canyon (n=18), an area known to be frequented by the Cuvier’s beaked whale (Bittau and Manconi, 2016). Specifically, in the Caprera Canyon area (Supplementary Figure S1) the sampling activity took place once a month from May up to October 2021, in 3 sites surmounting the canyon: an inshore location, named Station 1 (41°20’04.1“ N, 9°46’27.9” E) with a sea-bottom depth of approximately 550 m; a middle location, Station 2 (41°23’07.4“ N, 9°53’25.7” E) with a depth of approximately 750 m; and an offshore location, Station 3 (41°25’11.2“ N, 10°04’24.4” E) with an approximated depth of 1000 m. The choice of these 3 points was made to cover the entire environment of the canyon from east to west, sampling at different depths and at progressive distances from the coastline. Overall, a total of 18 samples were collected over 6 month-sampling activities. This group of samples was selected both in order to assess the efficiency of the developed species-specific molecular probe in detecting traces of the Cuvier’s beaked whale eDNA and, secondly, to assess its seasonal occurrence in the Caprera Canyon area. The second set of samples included “control samples” collected along a line crossing perpendicularly the Caprera Canyon (n=3), in order to verify whether Cuvier’s beaked whales do really favor waters above underwater canyons. Finally, the study was complemented with a third and larger sample set (n=32) collected in marine districts adjacent to the Caprera Canyon, (i.e. around Corsica, Sardinia and the Tuscan Archipelago) during the same period, within the project *Spot the Monk* (Valsecchi et al., 2023). All 53 samples were collected in 2021, from the 16^th^ of May until the 12^th^ of November and their full details are listed in Supplementary Table S1.

### 2.2 Sampling Activities and Seawater Filtration

All samples were collected from the most superficial layer of the sea (0–30 cm below sea level) from the research vessel. For each sample, a total of 12L of marine water were collected by pumping in a resistant Flexmet made sterile Bags-in-Box containers, following Valsecchi et al. (2021). Once the containers were filled, they were stored in a dark and fresh place to avoid an elevate exposition to heat and UV to minimize the degradation of DNA traces. All filtering activities were carried out within 12 hours from water collection, for the Caprera Canyon samples, while for the remaining samples the time elapsing between collection and filtration could reach 52 days (Valsecchi et al. 2023), with an average of 11.2 days. Each bag was divided into three 4-L aliquots, each filtered on a nitrocellulose filter with a porosity of 0.45 μm using the BioSart 100 filtration cylinders (Sartorius), resulting in filters A, B and C. For some of the samples collected outside the Caprera Canyon only one (A) or two (A and B) filters were obtained (Supplementary Table S1, see also Valsecchi et al. 2023). The water sample was forced to pass through the filter thanks to the negative pressure created by means of a vacuum pump (Fisherbrand FB70155, Fisher Scientific) applied to the water-collection vacuum flask.

After filtration the pored membranes were folded in two (filtrate side touching itself) inside an aluminum foil and accordingly labeled (sample id, location, date of sampling and filtration and filter number). Labelled filters were stored at −18°C before the DNA extraction at the University of Milano-Bicocca - MaRHE Center Lab.

### 2.3 Molecular Analyses

Environmental DNA was extracted using DNeasy PowerSoil Kit® (Qiagen), following the manufacturer’s protocol. Candidate regions for designing the Cuvier’s beaked whale specific primers were searched for within the mtDNA regions targeted by MarVer primers (12S-rDNA and 16S-rDNA), as this part of the mitogenome has proven to be highly polymorphic among vertebrates (Valsecchi et al., 2020). The candidate sets of primers were first tested on a panel of control tissue-extracted DNA templates (non eDNA) consisting of: *Ziphius cavirostris* DNA (positive control), and three mock templates containing of a mixture of a) fish DNAs (negative control), b) fish and cetaceans other than *Ziphius cavirostris* DNAs (negative control) and c) fish and cetaceans species DNAs including *Ziphius cavirostris* (positive control).

The 53 eDNA samples were screened using the best-performing set of newly designed primers for the detection of Cuvier’s beaked whale DNA traces, through Real Time (RT) quantitative PCR (qPCR), using an Applied Biosystem AB 7500. For each reaction were estimated the following parameters: the amplification efficiency (E), the Limit of Detection (LOD) and the Limit of Quantification (LOQ) (Klymus et al. 2020). To standardize the Ct (cycle threshold) we purified and isolated the amplicon extracted from *Ziphius cavirostris* tissue sample obtaining a control template with a concentration of 284 ng/μl and we used it to run a seven-fold serial dilution series to standardize the curve. For the amplification reaction we used: 5.0 μl SsoFast EvaGreen Supermix with Low ROX (Bio-Rad), 0.1 μl each [10 μM] primer solution, 2 μl eDNA template and 2.8 μl of Milli-Q water Q-PCR. The thermocycler profiles consisted of the following steps: 10 minutes at 95°C for the initial denaturation, followed by 40 cycles with denaturation at 95°C for 15s and 1 min of annealing-elongation at 52°C and final dissociation stage. According to the LOQ calculated for this locus, qPCR DNA detection outcomes were divided in three classes: 1) No signal; 2) Cuvier’s beaked whale eDNA detectable but not quantifiable (DBNQ); 3) positive quantifiable detection (PQD) of Cuvier’s beaked whale eDNA.

### 2.4 Data Analyses

Once obtained, the molecular data were divided into two groups, positive samples and negative samples, to investigate whether the incidence of positives was in any way related to the geographical characteristics of the sampled spots (i.e. depth of the seabed and distance from the coast), since the target species, being a deep-diver, has restraint requirements. Similarly, the two categories of results were investigated also to assess if the inhomogeneity in the processing times of the samples (namely time between sampling and filtering, that was immediate in the Caprera Canyon samples, while extremely variable and much longer in the remaining samples, collected through Citizen Science campaigns) could have affected the results.

## 3. Results

A set of primers (out of three tested) identified within the 16S-rDNA region showed the best amplification yield (strong specific band and no amplification in related taxa) in PCR tests. The primers’ pair was composed of the forward primer ZcaMV3F (5’CCCAAAAACTATAAATCTAAACCG3’), unique to the *Ziphius cavirostris* mitogenome (GenBank Accession Number NC_021435), and the reverse primer Ceto3R (5’TTGGATCAATAWGTGAT3’) conserved among most Cetaceans (Valsecchi et al. in prep.), amplifying in combination a 152bp amplicon.

In the standardization-curve qPCR run the amplification efficiency reached 86,075 % LOQ corresponded to Ct= 34.44, while the melting temperature (TM) of the targeted amplicon was of 80.8 °C. A total of 444 qPCR reactions were run on the 53 eDNA samples. All negative controls (run in triplicates in each qPCR run) did not produce any signal indicating presence of *Ziphius cavirostris* genetic traces, excluding the possibility of cross-sample contamination. The qPCR results were divided in the three categories: No detection, Positive and Quantifiable Detection (PQD) and Detectable But Not Quantifiable (DBNQ), according to the LOQ classification established in the protocol. Detections (both PQD and DBNQ) were considered reliable when amplified products were supported by the specific melting temperatures recorded (i.e. 80.8 +/-0.3) in the dissociation test.

DNA traces (PQD and DBNQ detections combined) of *Ziphius cavirostris* were detected in 41 of the 53 samples (77.4%), but in about half of these (n=26, 49.1%) the signal was intense enough to be quantifiable (PQD). The screening outcomes are summarized below, in Figure 4. Despite both PQD and DBNQ outcomes are indicative of the presence of Cuvier’s beaked whale DNA in the sampled water column, the latter denotes the presence of traces so diluted that they could be residues of signals released far away from the sampling point (probably tens of km apart). Since one of the purposes of this study is to identify fine-scale differences of presence-absence in sampling stations whose reciprocal distance does not exceed 30km (e.g. those of the Caprera Canyon), we have given more emphasis in the discussion to the PQD results, which identify stronger signals and therefore presumably released by the animal near the sampling point.

The geographical distribution of all 53 samples is mapped in Figure 2, which also highlights the samples that tested positive according to either of two criteria (PQD and DBNQ) described above. Figure 3 focuses on the spatiotemporal distribution of positives in the 21 (18 plus 3 controls) samples collected monthly from the surface overlooking the Caprera Canyon rim.

**Figure 2.**
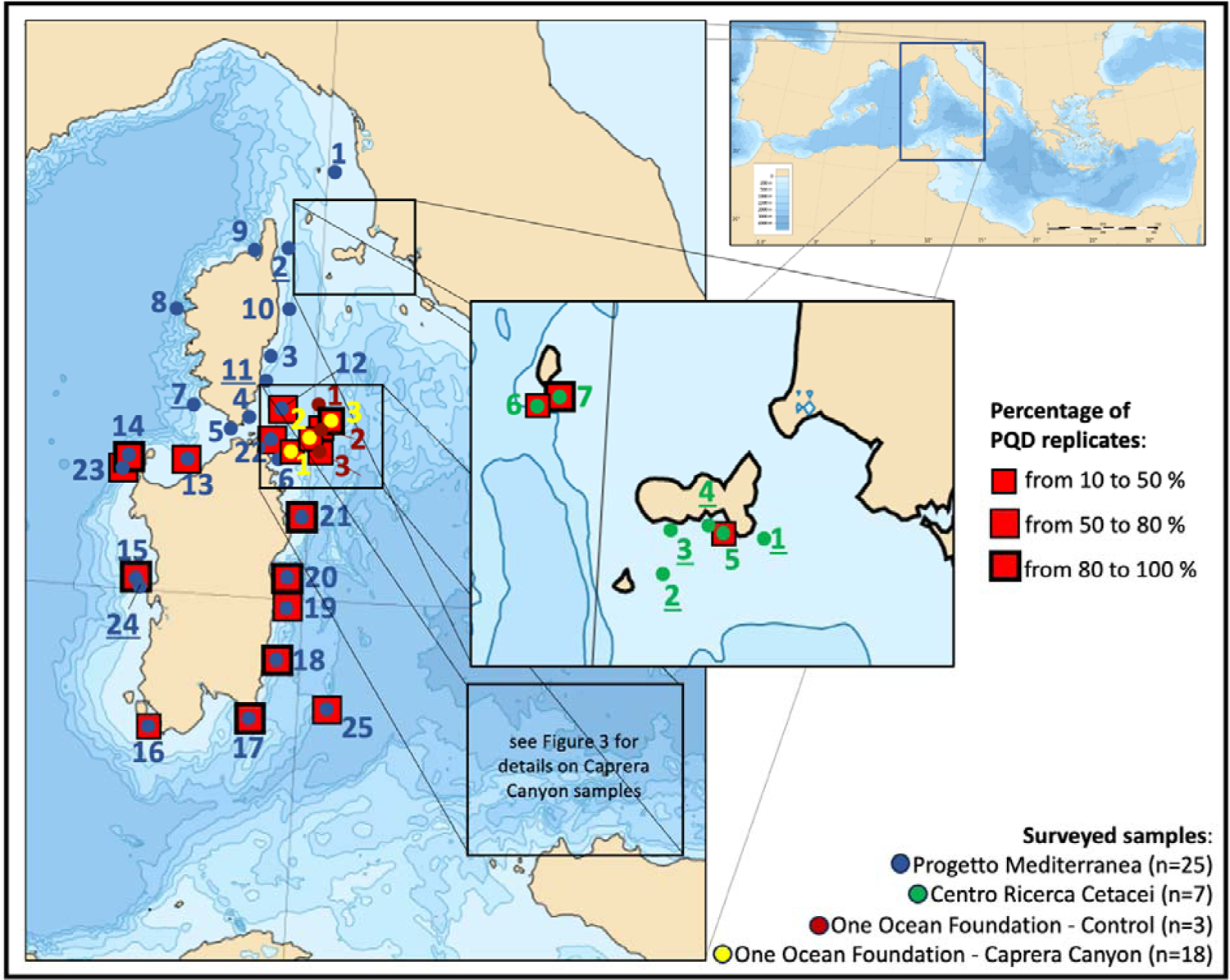
Maps showing the 53 points sampled in 2021 in the Northern Tyrrhenian Sea and Sardinan Channel (Central Mediterranean Sea, see top right map). Red squares surround samples that returned a positive and quantifiable detection (PQD) of *Ziphius cavirostris* eDNA traces. Larger squares depict a stronger signal, with more than 50% of the replicates returning a PQD detection (> 80% of replicates when squares are highlighted with a thicker outline). The numbers underlined indicate those points where a Detectable But Not Quantifiable (DBNQ) molecular signals referable to the Cuvier’s beaked whale were found. Maps downloaded from https://commons.wikimedia.org/wiki/File:Mediterranean_Sea_Bathymetry_map.svg.

**Figure 3.**
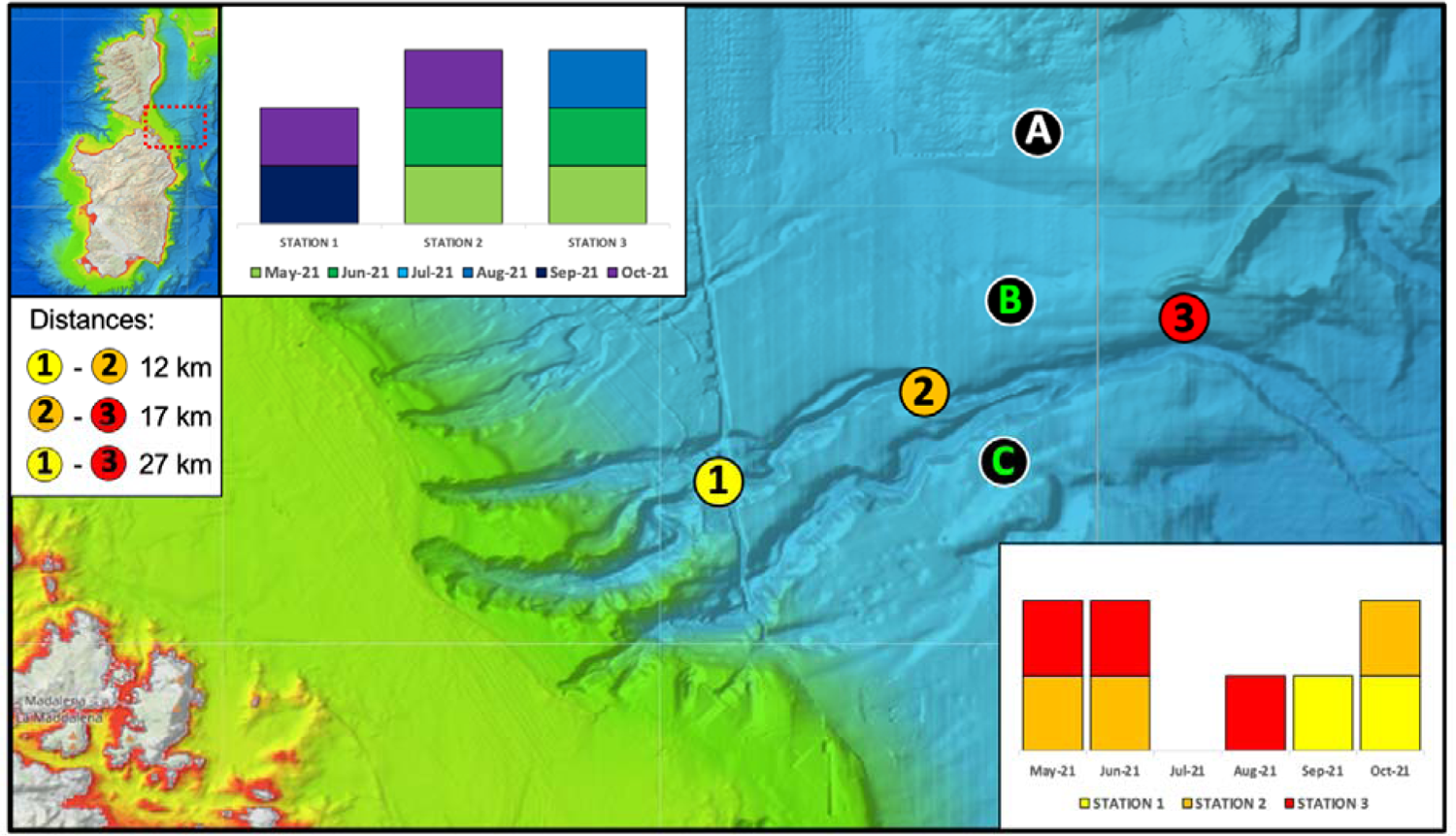
Caprera Canyon sampling sites’ map (area localised in the red dashed rectangular in the map in the left top corner). Coloured circles show the positioning of the three fix sampling stations sampled monthly from May to October 2021. The black circles indicate the 3 control samples surveyed in Nov 2021. The graph in the upper part of the figure shows the distribution of those samples returning a positive and quantifiable detection (PQD) in at least one of nine replicates, in the three fix stations over the six-month study period. While the graph below shows the monthly incidence of positive replicates in the three fix sampling stations. Only control samples B and C (shown in green) were positive to Cuvier’s beaked whale eDNA. Maps retrieved from https://www.emodnet-bathymetry.eu/.

In order to better appreciate seasonal differences, positives’ distribution was plotted in chronological order, highlighting a higher incidence of positives in the second half of the study period (Figure 4). The 26 PQD positives, accounting for roughly half (49.1%) of the total sample set, were found in samples collected, on average, in points with slightly higher bathymetry and further from the coast, compared to negative samples (as expected considering the pelagic habitat of the species, e.g. Arcangeli et al. 2016), however these differences were not significant (Supplementary Figure S2 A and B). Also, the long processing times of the Citizen Science samples did not seem to perturbate the incidence of positives (Supplementary Figure S2 C).

**Figure 4.**
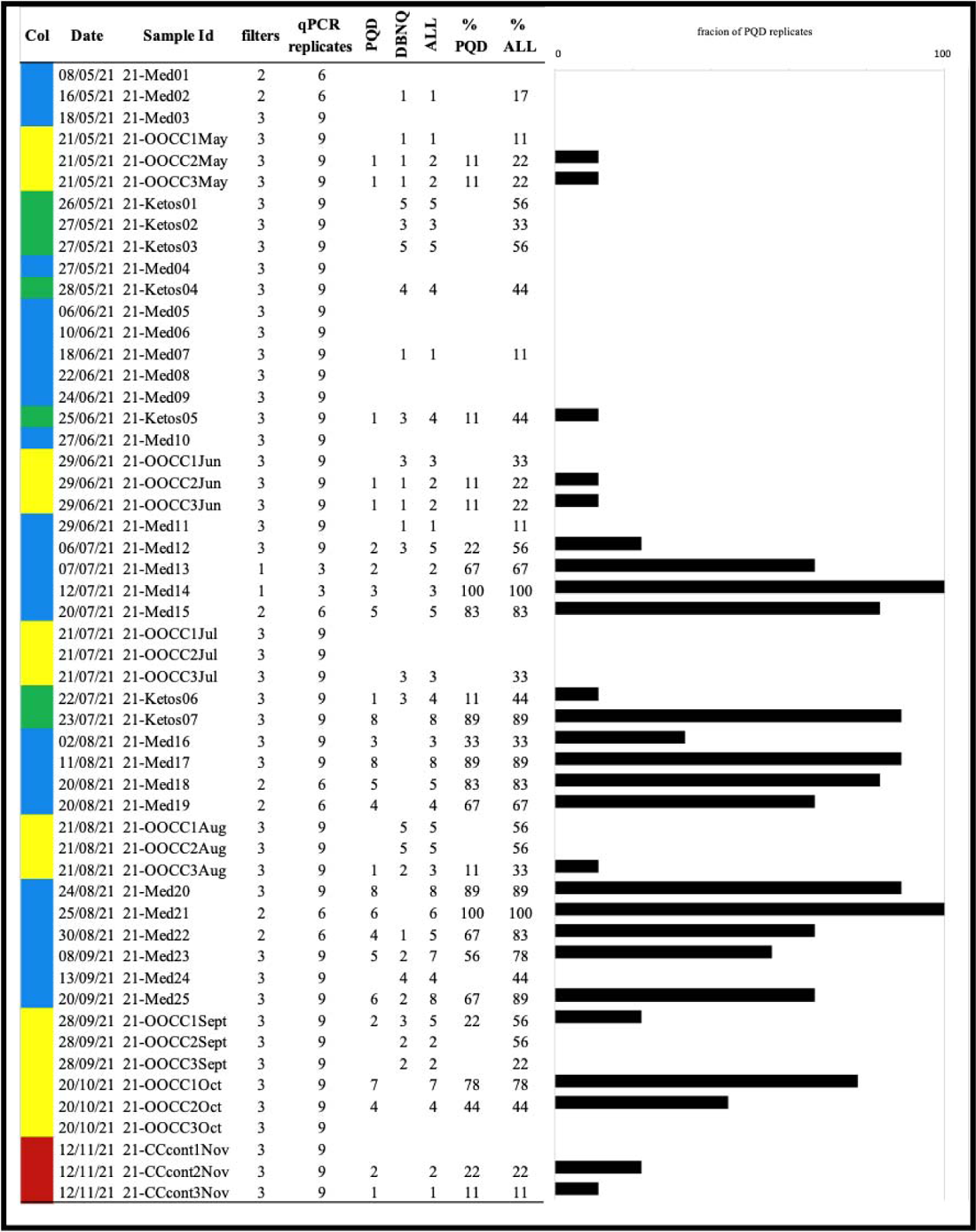
Summary of results of the qPCR assay for Cuvier’s beaked whale eDNA detection performed on the 53 samples, here ordered chronologically. From left to right are displayed: the color-code of samples as shown in Fig. 2; the date of sampling; the sample identification code; the number of filters obtained for each sample; the number of replicates (three per filter) tested in the qPCR assay. The next three columns (PQD, DBNQ, ALL) indicate the number of replicates in which either Positive and Quantifiable Detection (PQD), or Detectable But Not Quantifiable (DBNQ) or both were found, respectively. The final two columns indicate the percentage of replicates testing as positive for PQD (% PQD) or PQD and DBNQ (% ALL). Finally, the graph of the right depicts the distribution of PQD positive over the study period.

## 4. Discussion

In this study, a non-invasive, species-specific detection method, based on eDNA analysis, was successfully developed to monitor the presence of the endangered and elusive Cuvier’s beaked whale in the Canyon of Caprera, and in the wider Tyrrhenian Sea. Although DNA metabarcoding is more useful and cost-efficient when detecting greatest numbers of species at the same time (Gargan et al., 2022), cryptic species needs highly specific and sensitive approaches to be identified (Vörös et al., 2017). Compared to other cetaceans, the Cuvier’s beaked whale is one of the most difficult to study: these animals are able to dive at depth greater than 1 km (Schorr et al., 2014; Gnone et al., 2023) and for extended time period (1 hour, or more), and they also spend short periods at the surface (typically between 2 and 8 minutes) (Baird et al., 2006; Tyack et al., 2006; see also Figure 1). These traits make standard monitoring methods reasonably ineffective since for cetaceans that spend much of their life submerged, abundance estimates based on visual surveys become challenging and prone to underestimates. As a matter of fact, it has been assessed that only 40% of cetacean groups that are directly on the transect line would be detected in outstanding conditions during visual sighting surveys conducted from a large vessel (Barlow, 2015). Even so, our current knowledge on this and other cetacean species largely rely on the visual approach, which remains the most informative to date, especially if based on long-term data (Cañadas et al. 2018, Gnone et al. 2023). Moreover, when dealing with Cuvier’s beaked whales, traditional surveys that incorporate boat- and aerial-based surveys, as well as satellite-tracked tags, and offshore passive acoustic monitoring may result in very expensive monitoring programs. On the contrary, detection through eDNA does not require approaching the study species, reducing the probability of accidental harm, either through vessel collision or through a disruption in the species’ sonar navigation ability (Qu and Stewart, 2019). Collecting water samples for eDNA analysis also overcomes the weather condition constraints imposed to visual survey methods, allows nocturnal surveys and requires less technical expertise than accurate taxonomic proficiency in morphological identification (Qu and Stewart, 2019; Suarez-Bregua et al., 2022; Valentini et al., 2019).

In this context, the Caprera Canyon has represented a perfect case scenario for monitoring the presence of the Cuvier’s beaked whale based on eDNA sampling. Indeed, in this area sightings of the species, based on traditional boat-based visual monitoring carried out from 2011 to 2019, covered only 6.6% of effort time over the 9 years period (Dr Luca Bittau personal communication) due to dominant winds that determines strong weather events (Gerigny et al., 2011), limiting the possibility of visual surveys. Moreover, besides the rarity of outstanding weather conditions, the Canyon of Caprera extends offshore approximately 15-30 nautical miles from the north-eastern coast of Sardinia, requiring a considerable effort in term of both time, trained operators, and costs.

On the contrary, in this study, we have demonstrated that the eDNA approach has a good potential for detecting elusive species in open-ocean conditions and carrying out long term monitoring program. As a matter of fact, positive detection showed a constant presence of the Cuvier’s beaked whale in the Caprera Canyon (except in July) (Figure 3), highlighting the importance of this area for this species. Preliminary results based on visual surveys carried out in 2011-2013 have speculated this area as a favorable habitat for the Cuvier’s beaked whale (e.g. Bittau and Manconi, 2016; Gnone et al., 2023), and the results presented here, with data collected in recent year and with a continuous monitoring, can only support this hypothesis. In addition, the control stations used in this study (Figure 3) confirmed this result as well: the Cuvier’s beaked whale was detected only in stations B and C, which are located in proximity to the Canyon, but not in station A, located approximately 12 km north of it.

Within the Caprera Canyon area, this species was mainly determined at station 2 and 3, which are characterized by a bathymetry of approximately 700-1000 m. This is line with what reported in previous studies from the Mediterranean Sea: for instance, most sightings from the Ligurian Sea were located between 756 and 1389 m (Moulins et al., 2007). The Cuvier’s beaked whale is often associated with steep slope habitat and submarine canyon as its most common prey species in the Mediterranean are oceanic and meso- or bathypelagic cephalopods, inhabiting depths of approximately 1000 m (Azzellino et al., 2012; Blanco and Raga, 2000; Cañadas and Notarbartolo di Sciara, 2018; MacLeod, 2005). Therefore, the habitat distribution of the Cuvier’s beaked whale in the Caprera Canyon likely reflect that of its preys. Interestingly, in fall (September and October), the Cuvier’s beaked whale has been detected in the most inshore station (Figure 3). This shift to shallower bathymetries may be linked to the inshore movements of its prey. Indeed, several squid species are known to undertake inshore–offshore movements (e.g. Agus, 2015; Arkhipkin, 2000; Pierce et al., 2008) and the Cuvier’s beaked whales may follow their migration ending up in more inshore areas. Alternatively, these movements may reflect seasonal changes in the preference for its prey items (Azzellino et al., 2008).

Consistently with what reported for the Canyon of Caprera monitoring, the opportunistic samples collected in the Tyrrhenian Sea further highlights the importance of submarine canyons for this species. By looking at positive detections, especially those with a stronger signal (>80%, Figure 2 and 4), it is evident that the Cuvier’s beaked whale inhabits most of the underwater canyons of the Tyrrhenian Sea. As a matter of fact, stronger signals were reported in samples 21-Med13 and 21-Med14, collected in the Castelsardo Canyon area (northeastern Sardinia), in sample 21-Med15 gathered in proximity of Oristano Canyon, and in the four consecutive samples 21-Med18, 21-Med19, 21-Med20 and 21-Med21 collected in the area of Orosei, Gonone and Arbatax canyon systems in the same period (from the 20^th^ to the 25^th^ of August 2021), and finally sample 21-Med17, in the Simius Canyon (Figure 2). However, contrarily to the monitoring carried out at the Caprera Canyon, it is impossible to determine if in these Sardinian submarine canyon systems, the presence of the Cuvier’s beaked whale is seasonal, year-round, or only transient. Future monitoring should be carried out to further investigate this aspect.

Interestingly, while this species is widely distributed around Sardinian’s underwater canyon systems, it was never detected around Corsica Island (Figure 2 and 4). The absence of detections might not be surprising when considering the eastern part of the island as this lacks of submarine canyons, but appears peculiar when looking at sites where samples 21-Med07, 21-Med08, 21-Med09 were collected: they are all lying on massive submarine canyon systems on the western side of Corsica. However, if we ignore the morphological characteristics of the Corsican coast, which would suggest the presence of a habitat congenial to the species, our results do not differ from what was observed by Cañadas et al. (2018) in a study based on stacked visual data collected over a 27-year period: sightings were scarce in the water surrounding Corsica despite the intense search effort in the area (see Figure 1 in Cañadas et al., 2018). While considering that Corsican waters may lack of suitable habitats for this species sounds as an unrealistic hypothesis, the most immediate explanation for the lack of positive PQD detections in the Corsican samples is that those samples were collected too inshore; but so were the remaining opportunistic (Citizen Science) samples (mean distance from the coast 15.3km, with a median value of 5.5km), some of which were tested positive in Sardinia and the Tuscan archipelago. Thus, a further consideration would be the “seasonality” effect. Our data show that positive samples were scarce in the months of May and June, not only in Corsican waters (all Corse samples were collected in May and June), but also in other marine districts, for example in the Tuscan archipelago (Figure 4): here the only two PQD positive samples (21-Ketos06 and 21Ketos07) were collected towards the end of July, in close spatiotemporal proximity to each other (22nd and 23rd of July 2021, at 14.2km from each other) not far from Capraia island. Thus, an arguable question would be: is it possible that Cuvier’s beaked whales come closer to the coast in the late summer months possibly due to seasonal movement of their preys? This still remains unclear. What is certain is that, where we were able to sample systematically on a monthly basis (i.e. Caprera Canyon), the same trend was found: traces of *Ziphius*’s DNA were found in Station 1 (the most coastal one) only in the months of September and October. Inshore and offshore sampling should be carried out year-round in the western side of Corsica Island to elucidate the results presented in this study.

Another element that cannot be overlooked is linked to one of the major limitations inherent in the use of the molecular approach: marine eDNA is strongly influenced by the effect of marine currents, being drift away embedded in the water masses. For this reason, we opted to focus on the interpretation of strong signals (PQD), likely released not too far from the sampling point. Nevertheless, until the dispersion dynamics of marine eDNA are clarified and modeled, we cannot exclude that the traces identified through such a sensitive methodology do not in fact come from afar. However, although in the Central Tyrrhenian Sea visual data suggests the species being predominantly pelagic (e.g. Arcangeli et al. 2016), it cannot be ruled out the possibility of occasional nocturnal incursions towards more coastal waters.

## 5. Conclusion

The study presents the first attempt to monitor a deep-diving cetacean species by mean of eDNA surveys not associated to its sighting. The approach proved its potential as a non-invasive molecular monitoring not only for assessing the species presence but also its seasonal movements. Specifically, this study provides evidence of the regular presence of the Cuvier’s beaked whale, a threatened species, in the Canyon of Caprera, and, more widely, the importance of submarine canyons as elective habitats for this species, thus the pivotal priority to their conservation. As a matter of fact, our findings, in addition to previous studies that have been carried out in the area, indicate that the Caprera Canyon can be considered a hotspot area for the Cuvier’s beaked whale. As conservation efforts are increasingly focusing at preserving critical habitats, rather than selective species, the Caprera Canyon should be considered at both national and international levels for strong protection. Anthropogenic activities, including maritime traffic, fishing pressure, and noise and chemical pollution may represent a threat for cetaceans living off north–eastern Sardinia. In this regard, and in consideration that the very same area (samples in fact!) showed the elective use of the canyon also by other marine mammals, such as the monk seal (Valsecchi et al, 2023) the designation of an *Important Marine Mammal Area* (https://www.marinemammalhabitat.org/) would increase the protection of the overall ecosystem, within the overarching approach of systematic marine spatial planning, and may ultimately lead to effective legal protection, such as a Marine Protected Area.

Ethical approval is not relevant to this study, as all our samples consisted simply of marine water. Thus, all methods were carried out in accordance with relevant ethical guidelines and regulations.

## Acknowledgements

With regards to the sampling carried out at the Caprera Canyon, the authors and One Ocean Foundation’s team (www.1ocean.org/) would like to thank Mr Ruggero Ceriali and Mr Piergiorgio Purini for providing technical support during sampling collection. The Canyon of Caprera Project led by One Ocean Foundation is supported by Rolex as part of its Perpetual Planet Initiative.

We greatly thank Antonella Bruno and Valeria Catapano for laboratory logistic support. For samples’ collection we are grateful to the associations Progetto Mediterranea (https://progettomediterranea.com/) and Centro Ricerca Cetacei (www.centroricercacetacei.org/). We indeed thank Eleonora Moretta of CodeZero Digital Communication (www.codezerodigital.com) for creating the illustrations (visual abstract).

## Author contributions

BG coordinated Caprera Canyon samples’ collection/filtration, contributed to ms conceptualization, writing and editing; LC carried out the bench work and used these samples for her master thesis project under EV’s and BC’s supervision; PG provided laboratory facilities; BR provided logistic support and contributed in funding field work; EV developed the qPCR assays, designing the new primers and optimizing experimental conditions, provided most samples available from parallel projects, conceptualize the experimental design and main data analyses and sketched the original draft. All authors contributed to the manuscript’s improvement and agreed to its submission in the current version.

## SUPPLEMENTARY INFORMATION

**Figure S1.**
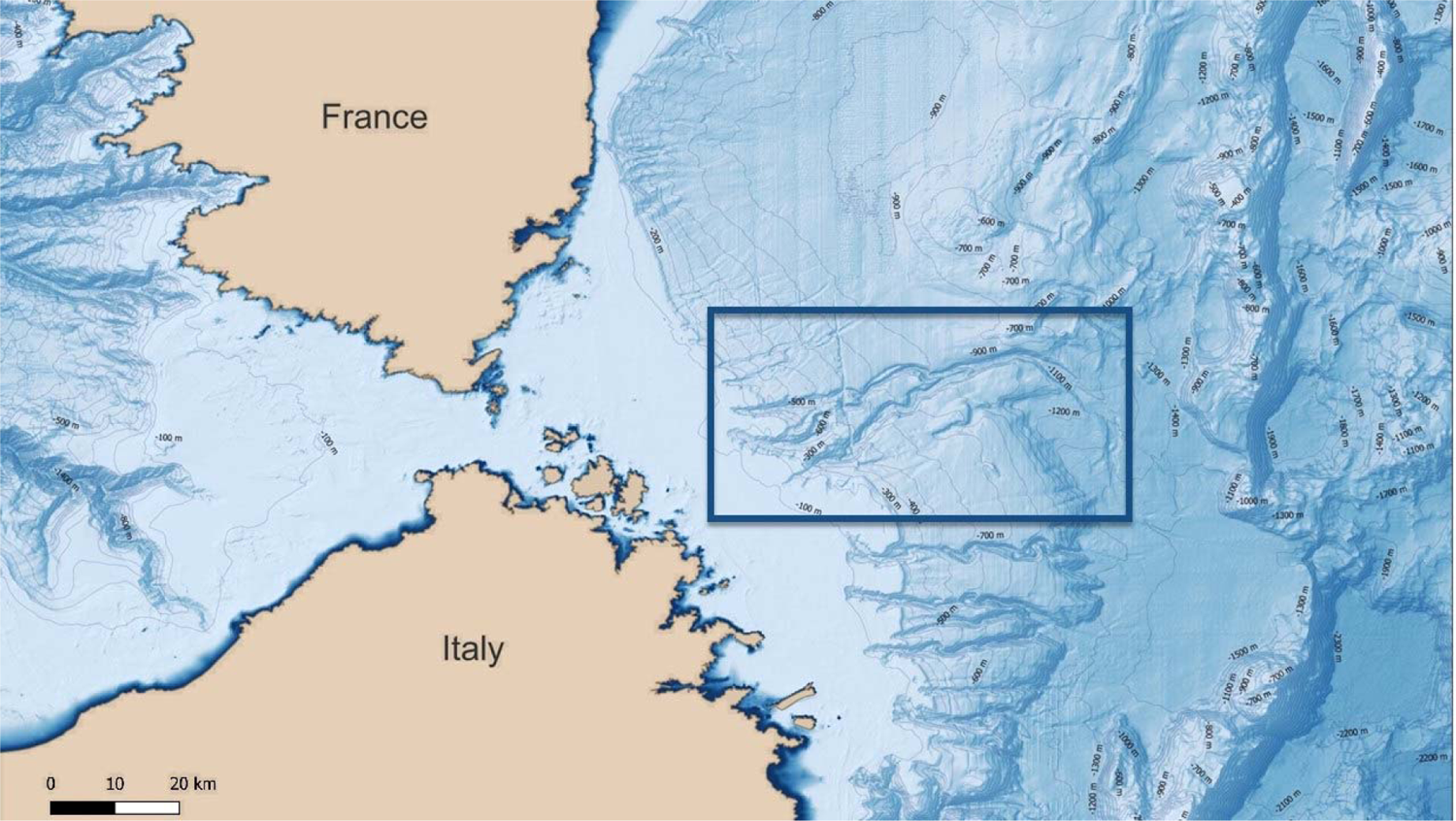
Bathymetric Map of the Caprera Canyon (extending from 41.27°N 9.70°E to 41.49°N 10.04°E approximatively) and adjacent waters. It is considered the largest system of submarine canyons in the northeast of Sardinia (central–western Tyrrhenian Sea).

**Figure S2.**
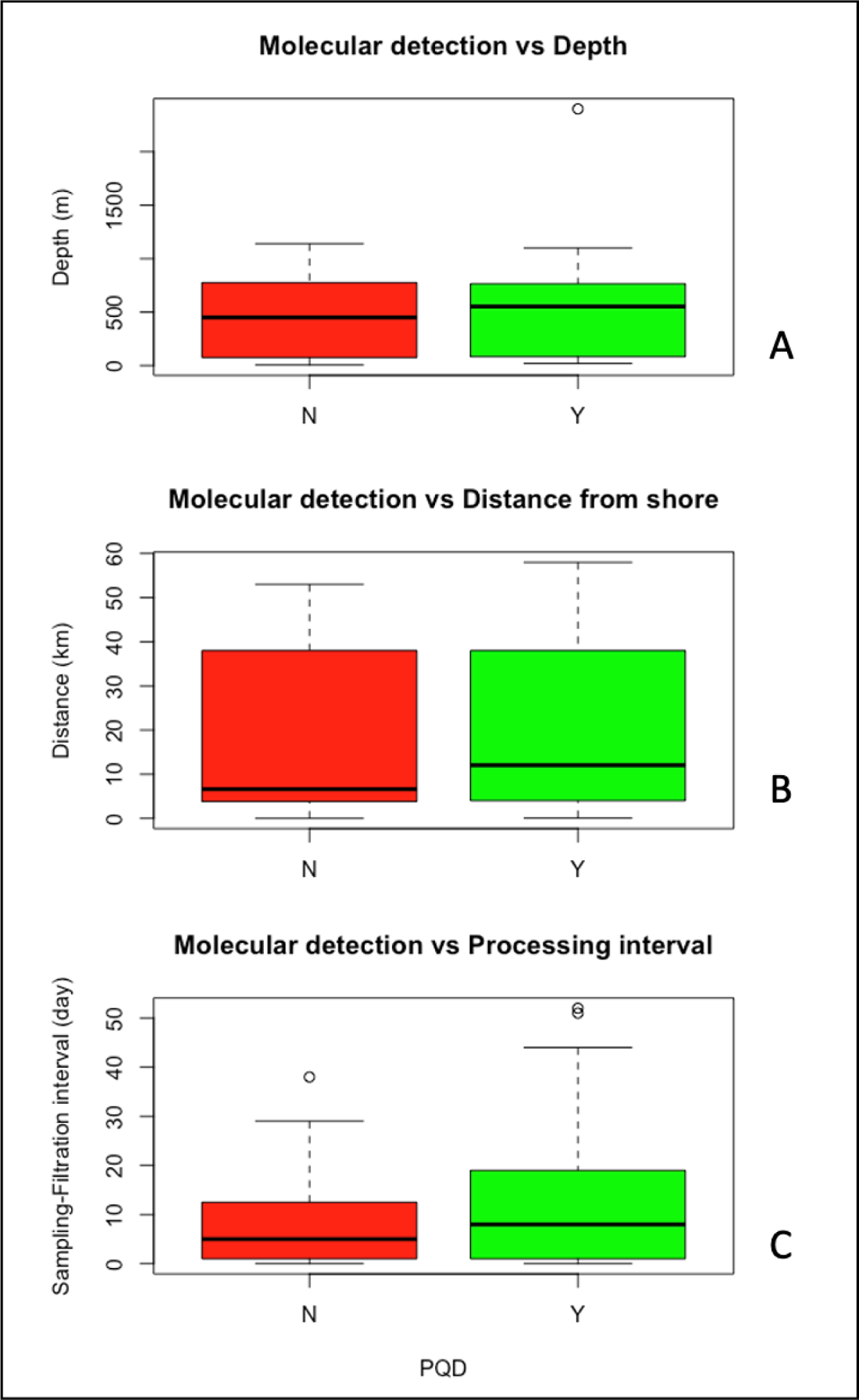
Molecular detection outcomes for *Ziphius cavirostris* eDNA traces in relation to: the bathymetry recorded at the sampling points (A); their distance from shore (B) and to the number of days between sampling and filtration of seawater bags (C). The box plot shows negative (N) and positive (Y) PQD detections. Central bar: median depth/distance from shore; Box: interquartile range; Whiskers: full range of data, excluding outliers; Circles: outlying values.

**Table S1.** List and characteristics of the 53 samples used in this survey, ordered by operator. The first column on the left (“Col”) indicates the color code of samples as shown in Figure 2. The Caprera Canyon samples are marked by the yellow color, the control samples taken at the intersection of the canyon are indicated in red, while the remaining samples taken in the adjacent marine districts are indicated in green and blue. The columns on the right-hand side of the geographic coordinates show respectively: the depth (recorded or retrieved) of the sea at the sampling point; its distance from the closest coastline (DisCo); how many days elapsed from sampling to filtration (“S→F”). The last two columns of the right summarize the molecular results, showing those samples returning, in at least one of the sample replicates, a positive detection (“Y”), for either only Positive Quantifiable Detections (PQD) or PQD and DBNQ (Detectable But Not Quantifiable) combined (“All”).

| Col | Date | Hour | Operator | Sample Id | Geocoordinates | Depth (m) | DisCo (km) | S→F (day) | PQD | ALL |
| --- | --- | --- | --- | --- | --- | --- | --- | --- | --- | --- |
| Yellow | 21/05/21 | 12:30 | One Ocean | 21-OOCC1May | 41°25'11.2"N 10°04'24.4"E | 1022 | 53 | 1 | N | Y |
| Yellow | 21/05/21 | 11:30 | One Ocean | 21-OOCC2May | 41°23'07.4"N 9°53'25.7"E | 765 | 38 | 0 | Y | Y |
| Yellow | 21/05/21 | 10:30 | One Ocean | 21-OOCC3May | 41°20'04.1"N 9°46'27.9"E | 555 | 27 | 0 | Y | Y |
| Yellow | 29/06/21 | 13:30 | One Ocean | 21-OOCC1Jun | 41°25'11.2"N 10°04'24.4"E | 1022 | 53 | 9 | N | Y |
| Yellow | 29/06/21 | 12:00 | One Ocean | 21-OOCC2Jun | 41°23'07.4"N 9°53'25.7"E | 765 | 38 | 0 | Y | Y |
| Yellow | 29/06/21 | 11:00 | One Ocean | 21-OOCC3Jun | 41°20'04.1"N 9°46'27.9"E | 555 | 27 | 0 | Y | Y |
| Yellow | 21/07/21 | 17:30 | One Ocean | 21-OOCC1Jul | 41°25'11.2"N 10°04'24.4"E | 1022 | 53 | 1 | N | N |
| Yellow | 21/07/21 | 15:15 | One Ocean | 21-OOCC2Jul | 41°23'07.4"N 9°53'25.7"E | 765 | 38 | 0 | N | N |
| Yellow | 21/07/21 | 13:45 | One Ocean | 21-OOCC3Jul | 41°20'04.1"N 9°46'27.9"E | 555 | 27 | 0 | N | Y |
| Yellow | 21/08/21 | 15:22 | One Ocean | 21-OOCC1Aug | 41°25'11.2"N 10°04'24.4"E | 1022 | 53 | 1 | N | Y |
| Yellow | 21/08/21 | 12:45 | One Ocean | 21-OOCC2Aug | 41°23'07.4"N 9°53'25.7"E | 765 | 38 | 0 | N | Y |
| Yellow | 21/08/21 | 11:00 | One Ocean | 21-OOCC3Aug | 41°20'04.1"N 9°46'27.9"E | 555 | 27 | 0 | Y | Y |
| Yellow | 28/09/21 | 15:22 | One Ocean | 21-OOCC1Sept | 41°25'11.2"N 10°04'24.4"E | 1022 | 53 | 1 | Y | Y |
| Yellow | 28/09/21 | 12:45 | One Ocean | 21-OOCC2Sept | 41°23'07.4"N 9°53'25.7"E | 765 | 38 | 1 | N | Y |
| Yellow | 28/09/21 | 11:00 | One Ocean | 21-OOCC3Sept | 41°20'04.1"N 9°46'27.9"E | 555 | 27 | 1 | N | Y |
| Yellow | 20/10/21 | 13:30 | One Ocean | 21-OOCC1Oct | 41°25'11.2"N 10°04'24.4"E | 1022 | 53 | 0 | Y | Y |
| Yellow | 20/10/21 | 12:15 | One Ocean | 21-OOCC2Oct | 41°23'07.4"N 9°53'25.7"E | 765 | 38 | 1 | Y | Y |
| Yellow | 20/10/21 | 11:20 | One Ocean | 21-OOCC3Oct | 41°20'04.1"N 9°46'27.9"E | 555 | 27 | 0 | N | N |
| Red | 12/11/21 | 07:00 | One Ocean | 21-CCcont1Nov | 41°30'46.4"N 9°58'29.9"E | 786 | 51 | 3 | N | N |
| Red | 12/11/21 | 11:00 | One Ocean | 21-CCcont2Nov | 41°26'00.0"N 9°57'00.0"E | 850 | 44 | 3 | Y | Y |
| Red | 12/11/21 | 16:00 | One Ocean | 21-CCcont3Nov | 41°21'01.8"N 9°56'51.6"E | 741 | 40 | 3 | Y | Y |
| Green | 26/05/21 | 10:40 | Centro Ricerca Cetacei | 21-Ketos01 | 42°42'18.0"N 10°27'42.0"E | 150 | 2.5 | 4 | N | Y |
| Green | 27/05/21 | 11:53 | Centro Ricerca Cetacei | 21-Ketos02 | 42°36'01.2"N 10°11'14.4"E | 75 | 7 | 3 | N | Y |
| Green | 27/05/21 | 22:37 | Centro Ricerca Cetacei | 21-Ketos03 | 42°43'30.6"N 10°09'29.4"E | 75 | 0.05 | 5 | N | Y |
| Green | 28/05/21 | 12:17 | Centro Ricerca Cetacei | 21-Ketos04 | 42°43'24.6"N 10°19'39.0"E | 75 | 1.4 | 5 | N | Y |
| Green | 25/06/21 | 19:05 | Centro Ricerca Cetacei | 21-Ketos05 | 42°43'04.4"N 10°22'32.6"E | 20 | 0.1 | 10 | Y | Y |
| Green | 22/07/21 | 11:35 | Centro Ricerca Cetacei | 21-Ketos06 | 42°57'47.6"N 9°43'53.3"E | 450 | 7.5 | 4 | Y | Y |
| Green | 23/07/21 | 09:19 | Centro Ricerca Cetacei | 21-Ketos07 | 42°59'09.3"N 9°54'11.8"E | 150 | 7.2 | 4 | Y | Y |
| Blue | 08/05/21 | 19:30 | Mediterranea | 21-Med01 | 43°13'17.7"N 9°58'51.7"E | 250 | 20 | 15 | N | N |
| Blue | 16/05/21 | 14:55 | Mediterranea | 21-Med02 | 42°41'43.3"N 9°27'07.7"E | 5 | 0 | 7 | N | Y |
| Blue | 18/05/21 | 15:10 | Mediterranea | 21-Med03 | 41°53'44.0"N 9°26'23.1"E | 40 | 3 | 5 | N | N |
| Blue | 27/05/21 | 18:15 | Mediterranea | 21-Med04 | 41°23'43.4"N 9°16'50.7"E | 73 | 2.8 | 38 | N | N |
| Blue | 06/06/21 | 17:00 | Mediterranea | 21-Med05 | 41°18'32.4"N 9°13'07.1"E | 73 | 5.6 | 29 | N | N |
| Blue | 10/06/21 | 18:10 | Mediterranea | 21-Med06 | 41°03'25.4"N 9°38'27.1"E | 71 | 5.1 | 25 | N | N |
| Blue | 18/06/21 | 11:20 | Mediterranea | 21-Med07 | 41°42'00.8"N 8°38'59.7"E | 450 | 5.3 | 17 | N | Y |
| Blue | 22/06/21 | 12:15 | Mediterranea | 21-Med08 | 42°18'13.4"N 8°32'50.8"E | 800 | 3.3 | 14 | N | N |
| Blue | 24/06/21 | 11:50 | Mediterranea | 21-Med09 | 42°49'02.6"N 9°14'22.1"E | 1140 | 5.9 | 12 | N | N |
| Blue | 27/06/21 | 16:40 | Mediterranea | 21-Med10 | 42°17'21.9"N 9°37'16.3"E | 95 | 5.1 | 9 | N | N |
| Blue | 29/06/21 | 11:05 | Mediterranea | 21-Med11 | 41°43'21.4"N 9°27'28.8"E | 420 | 4.3 | 7 | N | Y |
| Blue | 06/07/21 | 14:30 | Mediterranea | 21-Med12 | 41°13'55.8"N 9°37'06.0"E | 91 | 11.6 | 52 | Y | Y |
| Blue | 07/07/21 | 14:52 | Mediterranea | 21-Med13 | 41°08'20.8"N 8°42'26.3"E | 880 | 19.5 | 51 | Y | Y |
| Blue | 12/07/21 | 12:31 | Mediterranea | 21-Med14 | 40°58'25.7"N 8°03'44.0"E | 1100 | 10.9 | 44 | Y | Y |
| Blue | 20/07/21 | 18:00 | Mediterranea | 21-Med15 | 40°04'29.4"N 8°14'37.0"E | 140 | 12.5 | 37 | Y | Y |
| Blue | 02/08/21 | 10:45 | Mediterranea | 21-Med16 | 38°54'11.9"N 8°31'17.6"E | 82 | 7.4 | 24 | Y | Y |
| Blue | 11/08/21 | 12:45 | Mediterranea | 21-Med17 | 39°03'26.5"N 9°29'23.0"E | 550 | 4.9 | 16 | Y | Y |
| Blue | 20/08/21 | 08:50 | Mediterranea | 21-Med18 | 39°27'48.0"N 9°41'00.0"E | 62 | 3.1 | 19 | Y | Y |
| Blue | 20/08/21 | 16:50 | Mediterranea | 21-Med19 | 39°52'01.2"N 9°43'36.0"E | 110 | 2.9 | 19 | Y | Y |
| Blue | 24/08/21 | 10:30 | Mediterranea | 21-Med20 | 40°00'03.2"N 9°43'47.8"E | 73 | 2.5 | 15 | Y | Y |
| Blue | 25/08/21 | 15:30 | Mediterranea | 21-Med21 | 40°32'18.4"N 9°52'29.7"E | 63 | 4 | 14 | Y | Y |
| Blue | 30/08/21 | 10:55 | Mediterranea | 21-Med22 | 41°10'13.4"N 9°32'43.5"E | 70 | 2.2 | 27 | Y | Y |
| Blue | 08/09/21 | 12:42 | Mediterranea | 21-Med23 | 40°54'05.6"N 8°09'08.9"E | 80 | 3.5 | 18 | Y | Y |
| Blue | 13/09/21 | 12:35 | Mediterranea | 21-Med24 | 40°04'19.7"N 8°18'56.6"E | 71 | 6.6 | 13 | N | Y |
| Blue | 20/09/21 | 13:13 | Mediterranea | 21-Med25 | 39°02'30.6"N 10°14'20.9"E | 2400 | 58 | 6 | Y | Y |

## VISUAL ABSTRACT

With-text version

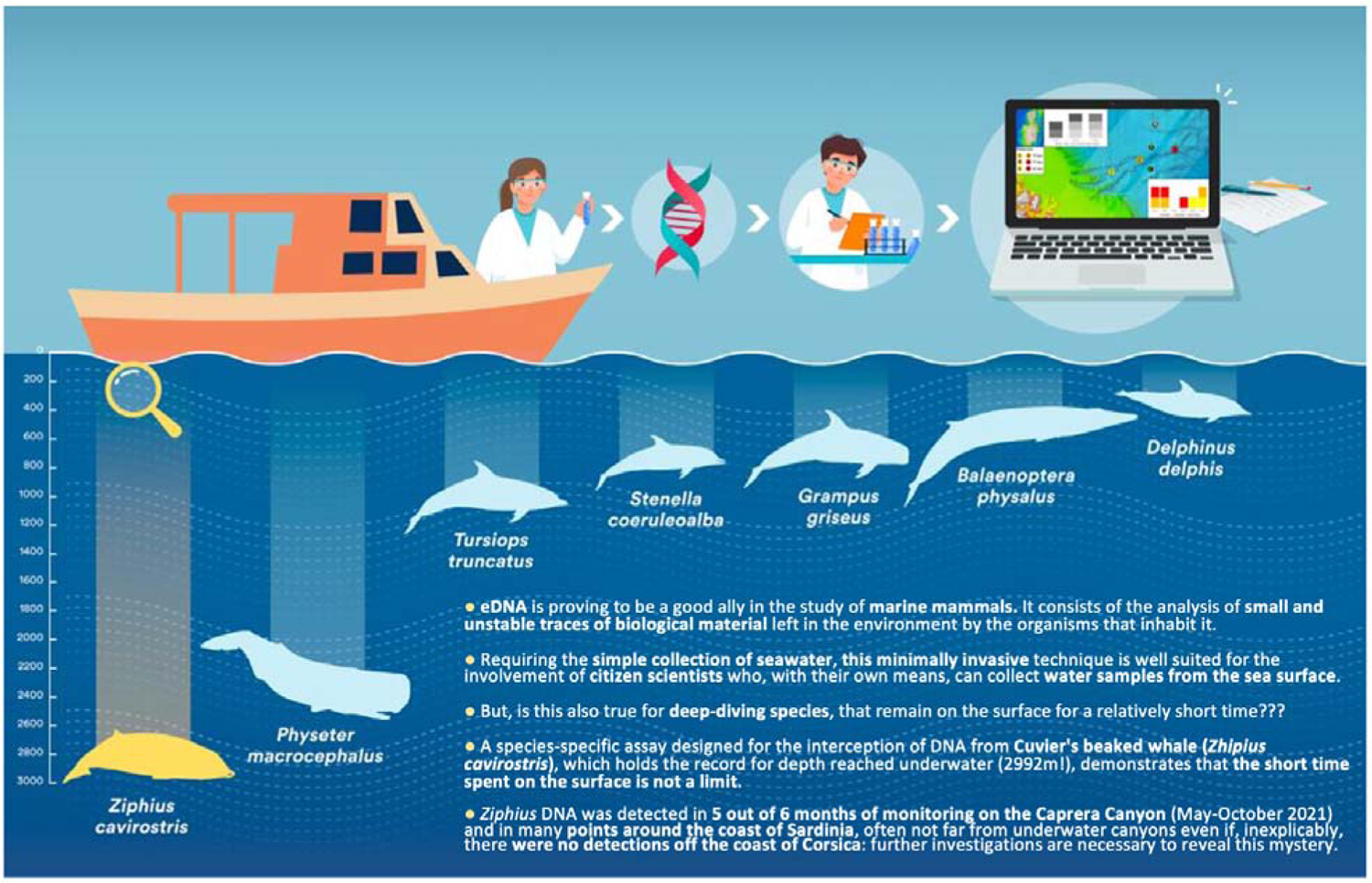

Without text version

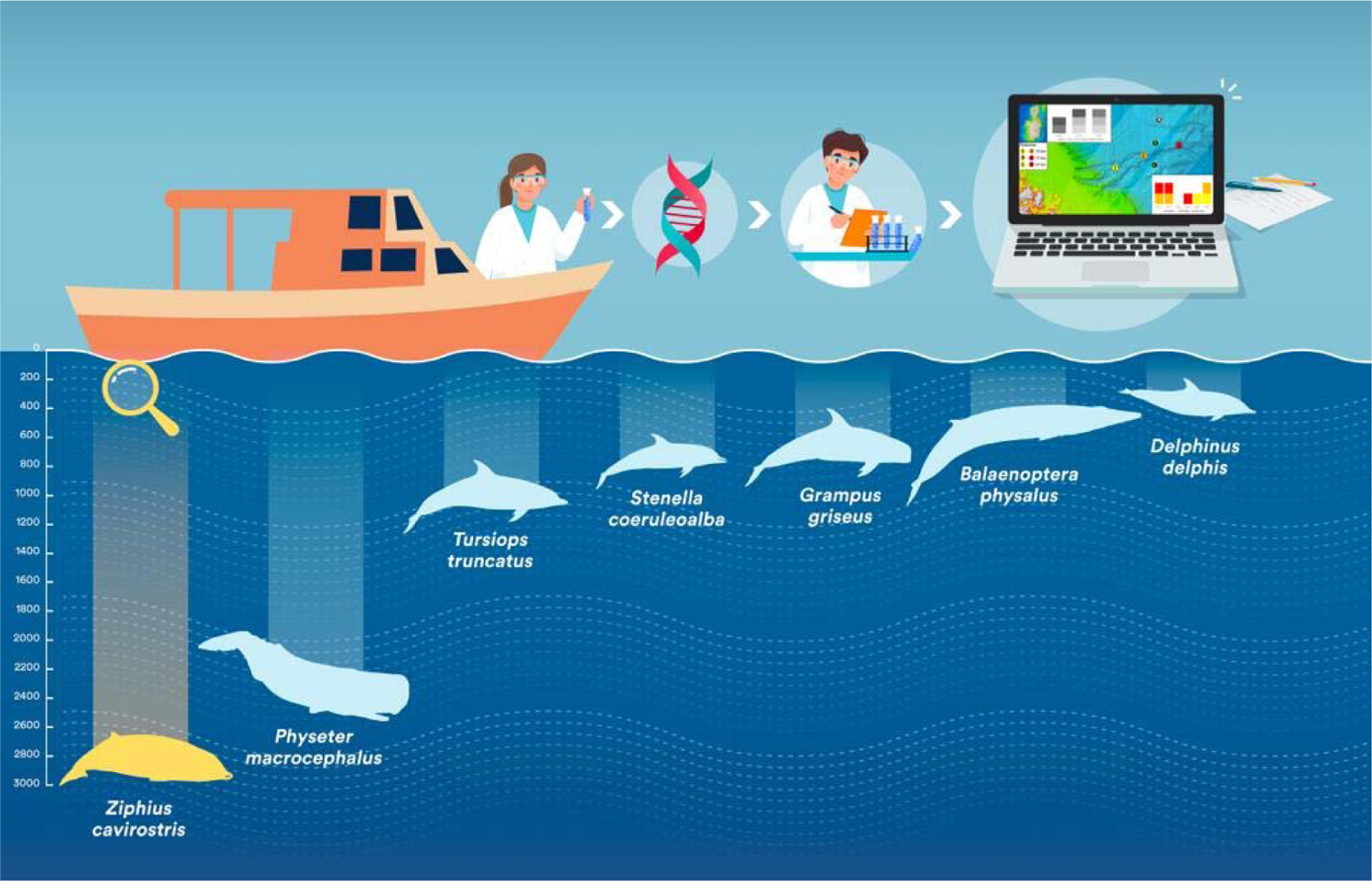

